# epialleleR: an R/Bioconductor package for sensitive allele-specific methylation analysis in NGS data

**DOI:** 10.1101/2022.06.30.498213

**Authors:** Oleksii Nikolaienko, Per Eystein Lønning, Stian Knappskog

## Abstract

Low-level mosaic methylation of the *BRCA1* gene promoter occurs in 5–8% of healthy individuals and is associated with a significantly elevated risk of breast and ovarian cancer. Similar events may also affect other tumour suppressor genes, potentially being a significant contributor to cancer burden. While this opens a new area for translational research, detection of low-level mosaic epigenetic events requires highly sensitive and robust methodology for methylation analysis. We here present epialleleR, a computational framework for sensitive detection, quantification and visualisation of mosaic epimutations in methylation sequencing data. Analysing simulated and real data sets, we provide in-depth assessments of epialleleR performance, and show that linkage to epihaplotype data is necessary to detect low-level methylation events. The epialleleR is freely available at https://github.com/BBCG/epialleleR and https://bioconductor.org/packages/epialleleR/ as an open source R/Bioconductor package.

## INTRODUCTION

Cancer is a major health threat and cause of death worldwide. While the minority of cases are due to highly penetrant germline pathogenic variants (inherited cancers), the majority are considered sporadic cancers with no known germline genetic component.

In addition to genetic aberrations like single-nucleotide variants, indels, copy number alterations and rearrangements, cancers are known to harbour epigenetic disturbances, leading to aberrant transcriptional up- and downregulation. Epigenetic aberrations may arise during different stages of carcinogenesis as somatic epimutations (mirroring somatic mutations), or *in utero* (affecting several germline layers) as constitutional normal tissue epimutations. Several studies in large cohorts (Lønning et al. 2018; Prajzendanc et al. 2020) have linked low-level mosaic epimutations to breast and/or ovarian cancer risk. Research and interest in this field, however, have been limited by the fact that all these studies were conducted on patients already diagnosed with their cancers, questioning whether normal tissue methylation in these patients may be a cancer-initiating event or a secondary effect of the disease itself. Recently we found frequent (occurring in >5% of healthy women) though low-level (down to 0.03% of affected alleles) mosaic methylation within the *BRCA1* gene promoter to be associated with a significantly elevated risk for subsequent high-grade ovarian as well as triple-negative breast cancer, in a large, population-based prospective cohort (Lønning et al. 2022). This finding raises a provoking question of whether similar low-level mosaic methylation may affect other tumour suppressor genes, and be associated with an elevated risk of other cancer forms as well. While this opens a new research area related to cancer risk, there are technical issues to account for, as the low frequency of such mosaic epimutations limits the amplitude of observed changes in methylation. Thus, to explore such hypotheses, there is a need for robust and sensitive methylation detection techniques.

Currently, the most widely used methods for DNA methylation profiling are BeadChip arrays (such as Illumina HumanMethylation450 or HumanMethylationEPIC) and a variety of methylation sequencing techniques (for details see (Sun and Zhu 2022)). These methods have different pros and cons: arrays allow genome-wide assessment at a reduced cost, while the sequencing provides additional information on haplotype specificity of DNA methylation. The typical bioinformatic workflows designed to analyse both types of data usually result in sets of beta values (ratio of a count of methylated cytosines to the total sum of methylated and unmethylated bases) for each genomic position covered (Krueger and Andrews 2011; Maksimovic et al. 2016; Fortin et al. 2017). While this approach is suitable for addressing large differences in DNA methylation profiles between two sets of samples (e.g., cases and controls), it lacks sensitivity for low-level mosaic methylation detection, as the detection is hindered by sometimes much more common biological variation (Youk et al. 2020; Gu et al. 2016) or technical artefacts (Kint et al. 2018; Stoler and Nekrutenko 2021). Moreover, the lack of haplotype linkage makes such analysis difficult in BeadChip array-based data sets, and therefore requires nontrivial approaches (Nikolaienko et al. 2022). In contrast, analysis of NGS-based data can provide much higher sensitivity when the base-resolution methylation data is combined with information on allelic belongingness (epihaplotype linkage).

Here, we present a computational framework for sensitive detection and quantification of low-frequency, mosaic epimutations in methylation sequencing data. The provided methods can be used for the discovery of low-frequency epialleles connected to disease risk (Lønning et al. 2022), as well as for purposes allowing less sensitivity, such as assessments related to treatment response (Kondrashova et al. 2018; Nesic et al. 2021), or to the development of treatment resistance (Hurley et al. 2021). Importantly, the framework also allows to connect DNA methylation status with potential underlying cis-factors, such as single-nucleotide variations or mutations within the immediate proximity.

## RESULTS

### epialleleR implementation

The presence of methylated *BRCA1* alleles in normal tissue (WBC) has been shown *qualitatively* for 5–8% of adult women (Lønning et al. 2018). However, the associated *quantitative* changes in DNA methylation at the level of individual CpGs are typically small (in most cases between 0.03 –1% (Lønning et al. 2022)), and therefore indistinguishable from the background methylation level due to inherent biological (potentially spurious single-base methylation events) and technical (sequencing errors) variance (Youk et al. 2020). Methylation statuses of neighbouring CpGs are often concordant (Huh et al. 2019), and such extended epigenetic changes are often associated with a gene expression silencing (Anastasiadi et al. 2018). Given the potential biological (gene inactivation) and clinical (cancer risk) importance of hypermethylated epialleles, we focused on quantification of methylation events that span over several CpGs, i.e. detection of epihaplotypes, accounting for both methylation status of individual CpGs within the sequence read as well as the average methylation level of the sequence read itself. This is possible in NGS-based data sets, while it is not in array-based data where methylation information of different CpGs cannot be connected to each other as in haplotype data. We also hypothesized that thresholding sequence reads by their average methylation level will reduce the effect of biological and technical variance and facilitate the detection of low-frequency hypermethylation events. Despite the trivial nature of the task and an increasing demand for such analyses, no suitable generic solution was publicly available. We therefore implemented a solution using R software environment for statistical computing (R Core Team 2021), a de facto standard for scientific data analysis. The implemented solution, epialleleR, loads methylation call strings and short sequence reads from supplied binary sequence alignment/map (BAM) file, optionally thresholds read pairs according to their methylation properties, and produces methylation reports for individual cytosines as well as genomic regions of interest (Fig. 1). During BAM loading, pairs of sequence reads and corresponding methylation call strings are merged according to Phred quality score values (i.e., base with the highest score is chosen) to preserve information of the highest quality. The optional thresholding defines a subpopulation of epialleles of interest and is based on the minimum number and the average methylation level of cytosines in various sequence contexts (e.g., CpG, CHG, or CHH). The thresholding parameters are fully adjustable to target desired population of epialleles; their default values (minimum 2 CpG sites, minimum average methylation beta value of 0.5 for CpG sites, maximum average methylation beta value of 0.1 for non-CpG sites) performed well in the study linking mosaic BRCA1 epimutations and cancer risk (Lønning et al. 2022) and were used here in all downstream analyses.

**Figure 1.**
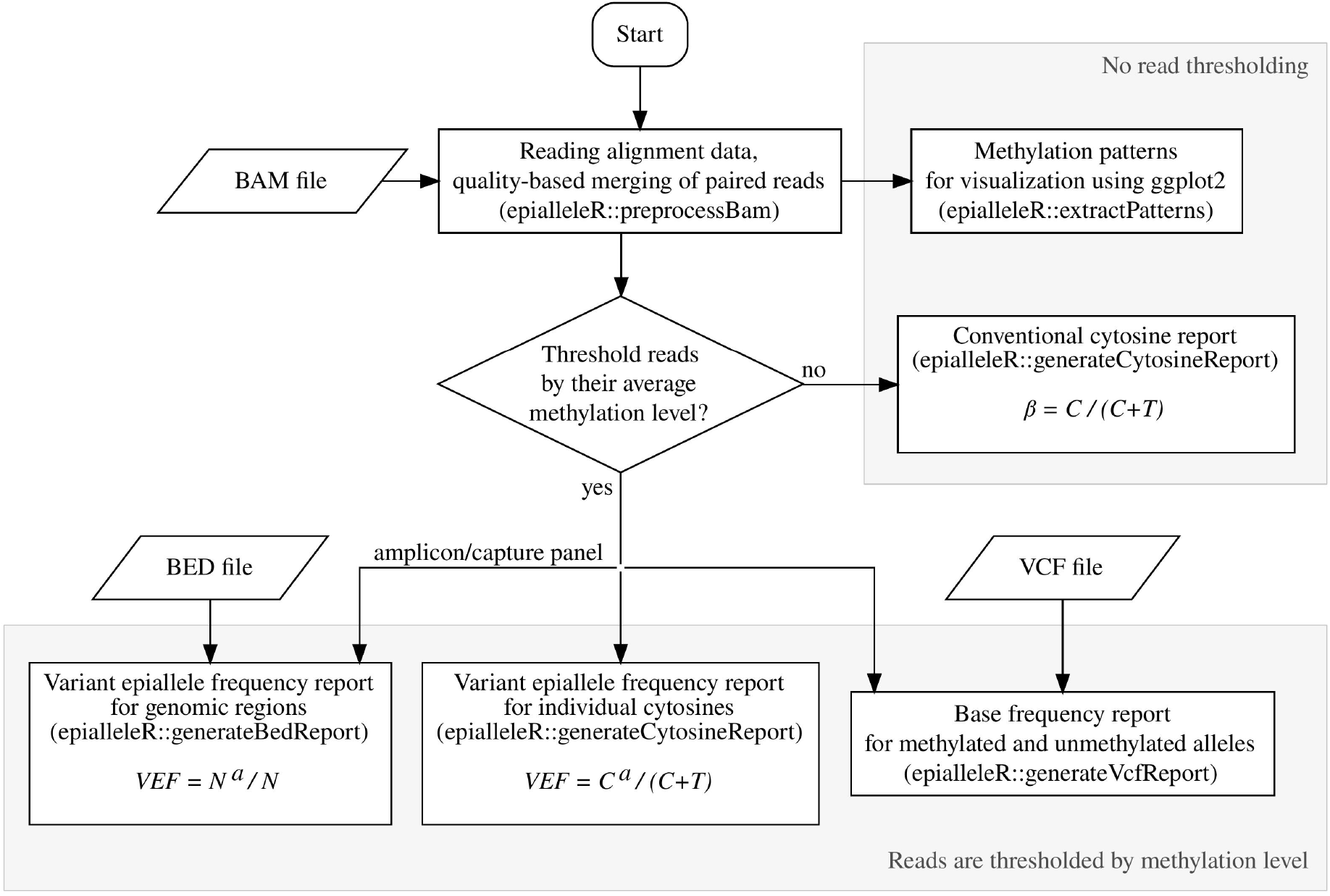
Flowchart of epialleleR package data processing steps. The formulas using to calculate conventional beta as well as VEF values are given in boxes. C and T, total number of cytosines and thymines at particular genomic position, respectively; C^a^, number of cytosines in read pairs passing a particular methylation threshold; N, total number of read pairs, mapped to a particular genomic region; N^a^, number of mapped read pairs, passing a particular methylation threshold.

The optional thresholding of sequence reads defines two modes of epialleleR function. Without thresholding, epialleleR produces conventional cytosine reports similar to the ones produced by other tools (e.g., Bismark (Krueger and Andrews 2011)). In this case, methylation beta value for every genomic location is computed as a ratio of a number of methylated cytosines to total number of methylated and unmethylated cytosines: *b = C / (C+T)*.

When read thresholding is performed (default mode of action), the level of methylation per every genomic position, denoted as a Variant Epiallele Frequency (VEF), is calculated as a ratio of a number of methylated cytosines in read pairs passing the threshold (*C*^*a*^) to total number of methylated and unmethylated cytosines in all read pairs: *VEF = C*^*a*^ */ (C+T)*. When the report is prepared at a level of extended genomic regions rather than individual bases, VEF equals to the ratio of a number of read pairs passing threshold (*N*^*a*^) to the total number of read pairs (*N*) overlapping the region of interest: *VEF = N*^*a*^ */ N*. The term “Variant Epiallele” here represents a group of epialleles (i.e., individual methylation patterns) with similar methylation properties that is defined by thresholding, therefore VEF effectively represents the frequency of this group of epialleles passing the threshold at the level of individual cytosines or extended genomic regions.

Methylation beta values (from conventional reporting) as well as VEF values (from default reporting mode with read thresholding) can be produced from any number of BAM files with no prior hypothesis, as long as experimental setup allows to call methylation on per-base level. Both of these values effectively represent methylation levels per genomic position and, as such, can be directly used further as an input for other bioinformatic tools including, but not limited to, differential methylation analysis tools.

To provide a comprehensive range of means for epiallele analysis, the package also offers methods allowing visualisation and characterisation of *all* individual epialleles (methylation patterns) in a sample (see Fig. 1 and Supplementary Figs for details). If required, extracted patterns can include other, non-cytosine bases of interest (e.g., single-nucleotide variations), which allows to connect methylation properties of epialleles with sequence features in proximity. During methylation pattern extraction, every epiallele is characterised by number of context sites and methylation level (average beta value) and is assigned with a unique identifier (Fowler-Noll-Vo FNV-1a non-cryptographic hash (Fowler et al.)) that solely depends on positions of included cytosine (and other optional) bases and their methylation states (or nucleotide symbols for optional bases), enabling not only to group epialleles by their methylation properties but also reliably and consistently track individual epialleles of high importance across different samples or even studies. The average beta values for all extracted patterns as well as patterns themselves can be explored to optimise thresholding parameters for a genomic region of interest.

Increasing scale and depth of methylation sequencing experiments impose a requirement on the speed of data processing. Therefore, all time-consuming subtasks were implemented using optimised C/C++ subroutines and, whenever possible, linked to HTSlib, unified C library for high-throughput sequencing data processing (Bonfield et al. 2021). The R package epialleleR is freely available at the Bioconductor package repository (http://bioconductor.org/packages/epialleleR/).

### Accuracy analyses

First, we sought to validate the accuracy of methylation reporting by epialleleR in its conventional mode (no read thresholding) as compared with three other commonly used tools for which read thresholding is not available: Bismark (Krueger and Andrews 2011), methylKit (Akalin et al. 2012) and Illumina DRAGEN Bio-IT Platform. For this purpose, we simulated large sets of paired-end bisulfite sequencing reads (2×151bp, 100 million read pairs covering human chromosome 19). In contrast to real datasets, simulated data allows to calculate “ground truth” methylation levels for unbiased comparison. Simulation parameters were selected to obtain exact methylation levels of 50% for cytosines in CpG context (n=2211240) and methylation level of approximately 0.25% (bisulfite conversion rate of ∼99.75%) for cytosines in CHG and CHH contexts (n=6593900 and 19210572, respectively). In addition to endogenous deamination events (Youk et al. 2020), bisulfite treatment-induced changes (Kint et al. 2018) and variation in conversion rates (Sun et al. 2021), sequencing itself can introduce errors that vary in range depending on assay type and sequencing technology (Stoler and Nekrutenko 2021). Therefore, we introduced variable level of artificial sequencing errors (0%, 0.1%, 0.3% or 0.6%) and evaluated their effect on the accuracy of reported methylation metrics, applying a selected set of methods (for comparison see Table 1). Analysis on exactly the same task (BAM file to cytosine report) revealed that reported values were close to their theoretical expectations for all methods, with epialleleR being the least affected by sequencing errors, i.e., maintaining the smallest deviance of reported versus expected methylation beta values for all samples with sequencing errors introduced, possibly owing to read quality-assisted merging of paired reads (Table 2, further details in Supplementary table 1).

**Table 1.** Selected characteristics of software/hardware solutions for methylation reporting.

| method | requires reference (genomic) sequence | removes overlaps within read pairs | outputs epiallele frequencies | processing speed, read pairs per second |
| --- | --- | --- | --- | --- |
| Bismark | yes | yes (trims read2) | no | 40–2,800 |
| methyKit | no | yes (trims read2) | no | 9,900–15,400 |
| DRAGEN | yes | yes (trims read2) | no | 2,000–183,000 |
| epialleleR | no | yes (base with the highest quality is chosen) | yes | 129,000–231,000 |

**Table 2.** Selected accuracy metrics (average beta values and their variance) of methylation reporting. Average beta values that are closest to the expected beta values and lowest variance values are shown in bold.

| sequencing error rate | method | CHH |  | CHG |  | CpG |  |
| --- | --- | --- | --- | --- | --- | --- | --- |
|  |  | mean | variance | mean | variance | mean | variance |
| 0.00% | DRAGEN | **0.002501** | 1.28E-05 | **0.002500** | 1.27E-05 | 0.499997 | 5.96E-07 |
|  | Bismark | 0.002501 | 1.34E-05 | 0.002500 | 1.33E-05 | **0.499997** | **5.92E-07** |
|  | methyKit | 0.002503 | **1.28E-05** | 0.002501 | **1.27E-05** | 0.499997 | 5.97E-07 |
|  | epialleleR | **0.002501** | 1.28E-05 | **0.002500** | 1.27E-05 | 0.499997 | 5.96E-07 |
| 0.10% | DRAGEN | 0.002661 | 1.37E-05 | 0.002662 | 1.36E-05 | 0.499835 | 1.73E-06 |
|  | Bismark | 0.002646 | 1.42E-05 | 0.002648 | 1.41E-05 | 0.499850 | 1.69E-06 |
|  | methyKit | 0.002662 | 1.37E-05 | 0.002664 | 1.36E-05 | 0.499835 | 1.75E-06 |
|  | epialleleR | **0.002624** | **1.35E-05** | **0.002626** | **1.34E-05** | **0.499872** | **1.54E-06** |
| 0.30% | DRAGEN | 0.002977 | 1.53E-05 | 0.002980 | 1.53E-05 | 0.499504 | 3.22E-06 |
|  | Bismark | 0.002928 | 1.57E-05 | 0.002931 | 1.57E-05 | 0.499554 | 3.03E-06 |
|  | methyKit | 0.002978 | 1.52E-05 | 0.002982 | 1.52E-05 | 0.499506 | 3.21E-06 |
|  | epialleleR | **0.002857** | **1.47E-05** | **0.002860** | **1.47E-05** | **0.499628** | **2.59E-06** |
| 0.60% | DRAGEN | 0.003498 | 1.79E-05 | 0.003506 | 1.78E-05 | 0.498942 | 1.29E-05 |
|  | Bismark | 0.003393 | 1.81E-05 | 0.003402 | 1.81E-05 | 0.499051 | 1.25E-05 |
|  | methyKit | 0.003497 | 1.78E-05 | 0.003501 | 1.78E-05 | 0.498952 | 1.28E-05 |
|  | epialleleR | **0.003237** | **1.66E-05** | **0.003241** | **1.65E-05** | **0.499222** | **1.14E-05** |

### Sensitivity analyses

Concordantly methylated alleles (genomic regions with spatially extended / haplotype-specific methylation) are thought to possess high biological importance (Lønning et al. 2022; Hurley et al. 2021). Spontaneous 5-methyl cytosine (5mC) deamination, sequencing errors, as well as genuine single-nucleotide methylation/demethylation events affect observed background methylation level and can therefore hinder the detection of low-frequency hyper- or hypomethylated alleles. Differences in experimental conditions provide an additional level of variability which can sometimes be tackled by normalisation during postprocessing (Aryee et al. 2014). In contrast to the DNA methylation analysis using BeadChip arrays (such as Illumina HumanMethylation450 and HumanMethylationEPIC) which report average methylation values at the level of individual cytosines only, next-generation sequencing provides an additional data dimension by linking methylation levels of individual nucleotides within a genomic region covered by a sequencing read (epihaplotypes). However, this information is lost when methylation is assessed and reported without accounting for its allelic distribution. To evaluate the sensitivity of detection for low-frequency monoallelic hypermethylation events in next-generation sequencing data, we simulated an extended set of samples using real, amplicon-based bisulfite sequencing data for human WBC (n=10 with almost no hypermethylated alleles, as described in Materials and Methods) and fully methylated control DNA samples. Combining real WBC DNA bisulfite sequencing data allowed to introduce sample-to-sample variability although maintaining biologically relevant background methylation levels across sequenced regions, while admixing fully methylated reads simulated low-frequency, concordant methylation events. The amplicons used, covered promoter regions of the tumour suppressors *MLH1, CDKN2A, MGMT, CDH1*, and *BRCA1*. The distributions of per-read beta values (Supplementary Fig. 1) and methylation patterns (Supplementary Fig. 2) of admixed samples show the expected abundance of hypermethylated (average b≥0.5) alleles and confirm their high similarity to the real samples (Supplementary Figs 3 and 4). Conventional cytosine reports (no read thresholding) as well as VEF reports (with read thresholding) were prepared and used for unsupervised clustering of samples and differentially methylated region (DMR) discovery. Despite quite low overall methylation level of amplified regions (average beta value of 0.014, median of 0.005; Fig. 2A), t-SNE analysis based on beta values was not able to discriminate between samples with 0.01%, 0.03%, 0.10%, 0.30% of methylated reads, or no methylated reads added (Fig. 2B, left panel). On the other hand, VEF value-based t-SNE analysis resulted in spatially well-separated clusters that corresponded to each level of admixed methylated reads (Fig. 2B, right panel). Intergroup DMR discovery based on beta values (Fig. 2C, left panel) showed fewer number of regions found as well as higher associated false discovery rate (FDR), while discovery based on VEF values resulted in all five possible regions identified for all possible intergroup comparisons as well as generally lower associated FDR. When each sample with admixed methylated reads was compared against the group of samples without admixed methylated reads, recall metrics for differential (by DMRcate (Peters et al. 2021); Fig. 2D) or aberrant (by ramr (Nikolaienko et al. 2022); Fig. 2E) methylation analysis were notably higher for analyses based on VEF values (Fig. 2D–E, right panels) in comparison with analyses based on beta values (Fig. 2D–E, left panels). This shows that VEF values are more valuable for detection and analysis of low-frequency (≤1%) hypermethylation events than methylation beta values.

**Figure 2.**
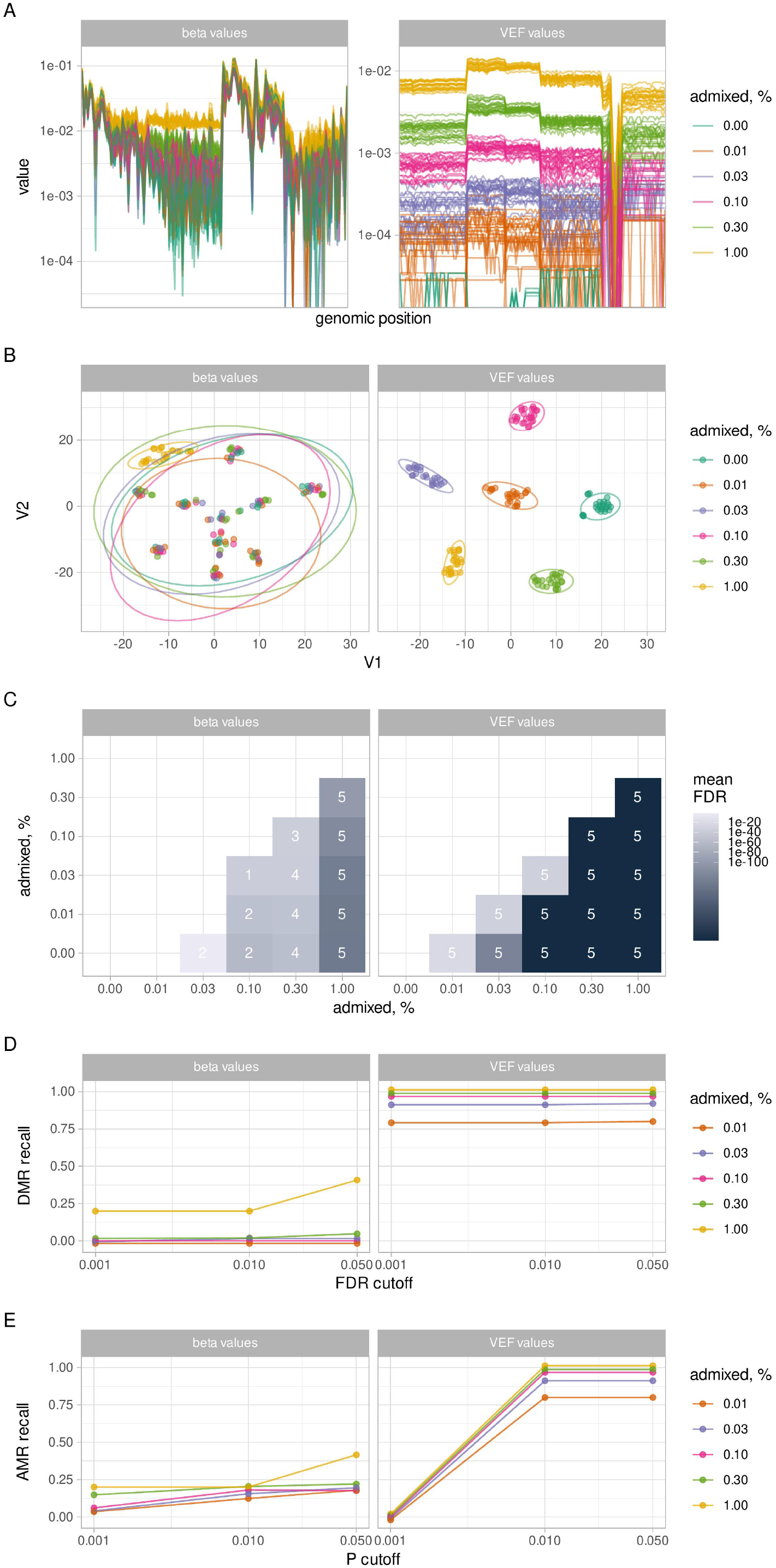
(A) Line plots of beta (left panel) and VEF (right panel) values for individual samples, colour-coded and grouped according to the amount of admixed methylated reads. (B) Embedding plots for t-SNE analysis using beta (left panel) and VEF (right panel) values. Ellipses represent 95% confidence levels. (C) Heatmap of mean false discovery rate for differentially methylated regions (DMRs) identified by DMRcate. Labels indicate the number of DMRs found (of total of five regions possible). (D) Recall rate for DMR identification using DMRcate for varying FDR cutoffs. (E) Recall rate for aberrantly methylated regions identification using ramr for varying p value cutoffs.

BeadChip arrays, such as Illumina HumanMethylationEPIC, is another widely used, amplification-free method to assess genome-wide DNA methylation for a reduced cost. In order to directly compare the sensitivities of targeted NGS and of the BeadChip arrays for the detection of low-frequency DNA methylation events, we employed both of the methods to analyse small set of samples (n=8) carrying low-frequency methylation in at least one of the assayed regions (promoter regions of *MLH1, CDKN2A, MGMT, CDH1*, and *BRCA1*). Sample distributions of per-read beta values (Supplementary Fig. 3) and methylation patterns (Supplementary Fig. 4) show that these samples indeed contain varying frequencies of hypermethylated (average b≥0.5) alleles. For unbiased comparison, we limited the corresponding data sets to the CpGs assayed and sufficiently covered by both techniques. Analysis revealed that VEF values of samples with many hypermethylated alleles (e.g., A26 and A45 for *BRCA1*; as apparent from Fig. 3A) differ significantly (Fig. 3B) from VEF values of samples with only a few or no hypermethylated alleles (e.g., A02 or A05 for *BRCA1*; Fig. 3A). When VEF values were used for identification of aberrantly or differentially methylated regions by ramr (Nikolaienko et al. 2022) or DMRcate (Peters et al. 2021), respectively, the significant regions found correlated well with the notable presence of hypermethylated alleles. Of note, slightly inferior performance of DMRcate is probably due to the fact that for some of the genomic regions too many samples in this subset simultaneously contained hypermethylated epialleles. When DMRcate was used for the same purpose on an extended set of sequenced samples (n=18, containing n=10 samples characterised by the absence of hypermethylated alleles that were used to create admixed sample set), its performance in identification of hypermethylated epiallele-containing samples was higher (Supplementary Fig. 5).

**Figure 3.**
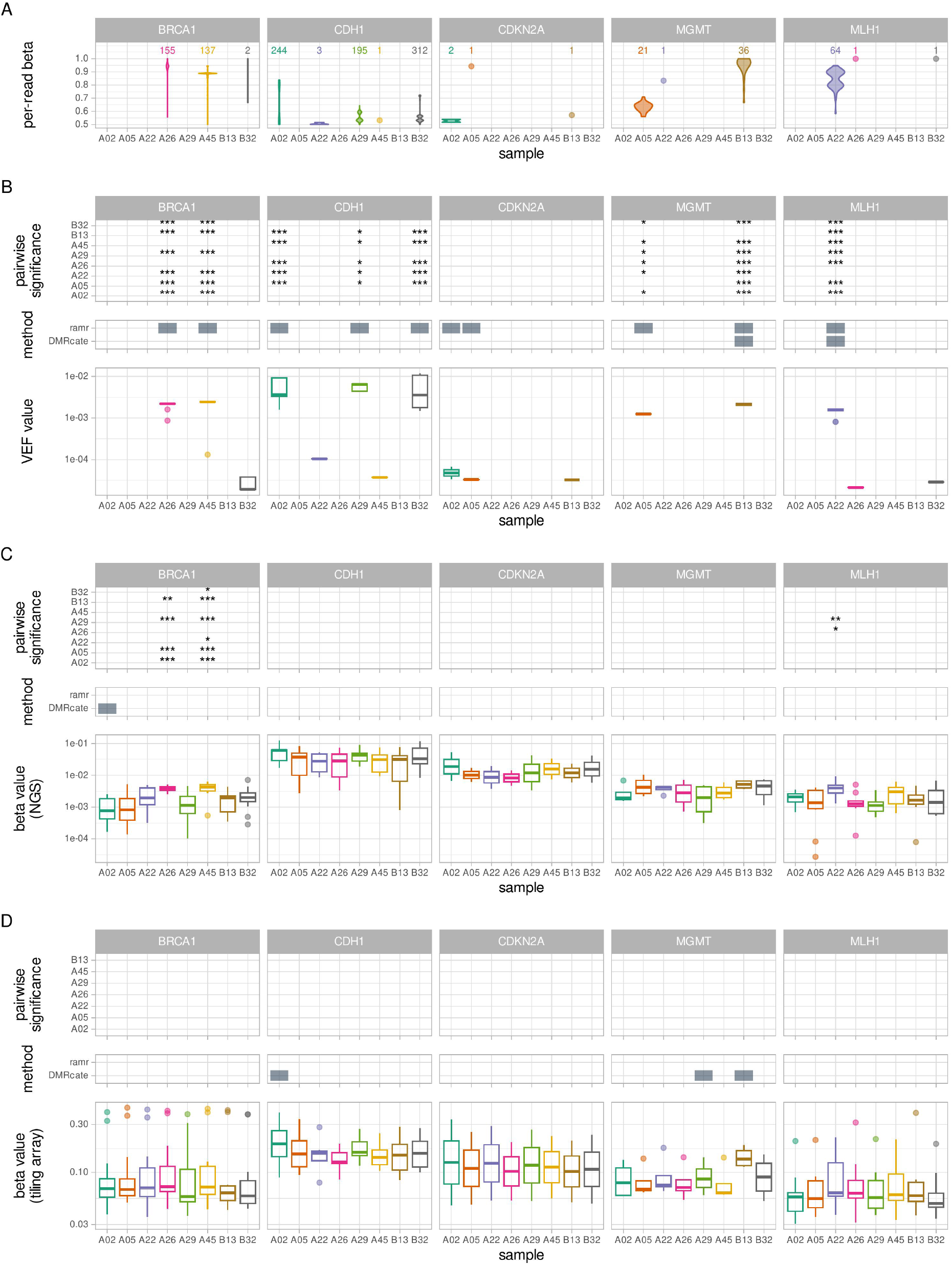
(A) Distribution of per-read beta values for NGS read pairs covering CpGs that are common for NGS and BeadChip array. For clarity, only the reads with average beta of at least 0.5 (i.e., representing hypermethylated epialleles) are included. Single observations are shown as dots, number of observations is given above. Complete density plots are provided in the Supplementary Fig. 3. Corresponding methylation patterns are provided in the Supplementary Fig. 4. (B) Lower panel: box plots of NGS-derived VEF values for individual CpGs; middle panel: significant aberrantly or differentially methylated regions identified by ramr or DMRcate, respectively, based on VEF values; upper panel: significance levels from pairwise comparison of VEF values. (C) Lower panel: box plots of NGS-derived beta values for individual CpGs; middle panel: significant aberrantly or differentially methylated regions identified by ramr or DMRcate, respectively, based on NGS-derived beta values; upper panel: significance levels from pairwise comparison of NGS-derived beta values. (D) Lower panel: box plots of BeadChip array-derived beta values for individual CpGs; middle panel: significant aberrantly or differentially methylated regions identified by ramr or DMRcate, respectively, based on BeadChip array-derived beta values; upper panel: significance levels from pairwise comparison of BeadChip array-derived beta values. (B–D) The lower and upper hinges of boxes correspond to the first (Q1) and third (Q_3_) quartiles; the bar in the middle correspond to the median value; the upper and lower whisker extend to Q_3_+1.5*IQR and Q_1_-1.5*IQR, respectively, while the values outside this range (outliers) are plotted as dots. Zero values are not plotted. *** p<0.001, ** p<0.01, * p<0.05, blank p≥0.05.

In contrast, only a few significant differences remained when NGS beta values were used for sample comparison (Fig. 3C), while pairwise comparisons based on BeadChip array beta values did not reveal any significant differences between samples (Fig. 3D). The search for aberrantly or differentially methylated regions using either NGS or array beta values did not result in identification of such regions in relevant (according to methylation patterns or beta value densities) samples. Generally higher beta values of BeadChip array as compared to NGS beta values likely mask subtle changes in methylation caused by the presence of infrequent hypermethylated alleles and hinder the detection of differences between samples.

Several scores to describe and quantify variability in DNA methylation in sequencing reads (within-sample heterogeneity, WSH) have been proposed (Scherer et al. 2020). In order to assess WSH, we evaluated the difference in combinatorial entropy between each pair of samples using methclone (Li et al. 2014) (Supplementary Fig. 6A). The largest (by absolute value) reported difference in combinatorial entropy of -2.59 between any pair of samples confirms a high similarity between sample methylation profiles, of note, being much smaller than cutoffs for epiallele shifts between samples analysed in (Scherer et al. 2020) (−60 and lower). Further, we also calculated four additional heterogeneity scores: combinatorial entropy, epipolymorphism, fraction of discordant read pairs (FDRP) and proportion of discordant reads (PDR). The scores themselves (Supplementary Fig.6B) and the levels of score-based pairwise significance between samples (Supplementary Fig. 6C), are not generally consistent with fractions of hypermethylated (average b≥0.5) alleles (Fig. 3A and Supplementary Fig. 3) or VEF values (Fig. 3B): e.g., samples A26 and A45 have a notable fraction of hypermethylated reads in *BRCA1* promoter region compared to other samples, although it is not reflected at the level of WSH scores. Importantly, WSH scores produced cannot be directly used as an input for DMR analysis tools, which are commonly employed to characterise exact differences in methylation between samples.

It is known that DNA methylation profiles of blood samples depend on the varying contribution of individual blood cell types (Reinius et al. 2012; Bakulski et al. 2016). While we cannot exclude that hypermethylated alleles present in the samples analysed here originate from a particular blood cell type, low-level, mosaic methylation of at least *BRCA1* was previously shown to be independent of blood subfraction composition (Lønning et al. 2018). Of note, only one CpG (cg05785947 in *CDH1*) out of 37 used in NGS vs BeadChip array comparison here, was found to be significantly differentially methylated between blood cell types of healthy males; and none of CpGs were significantly differentially methylated between blood cell types of newborns.

### Processing speed analyses

Methylation sequencing data produced by contemporary techniques varies in scale and depth and may contain several thousands to billions of single or paired-end reads. To analyse them efficiently, computational methods must be scalable and fast enough for as large as possible range of sample counts or data file sizes. Unfortunately, many academic tools use computationally complex algorithms that do not scale to contemporary tasks. We compared data processing speed for epialleleR versus methylKit, Bismark, and DRAGEN Bio-IT Platform, performing exactly the same task (BAM file to cytosine report) of methylation reporting across input data coming from various assays: amplicon-based (n=10 samples with a depth of coverage of ∼20,000x), genome-wide capture-based (n=10 with a depth of coverage of ∼60x and n=3 with a depth of coverage of ∼1000x) or whole-genome bisulfite sequencing (WGBS, n=6 with a depth of coverage of ∼60x). The obtained results confirm very efficient implementation of epialleleR and its suitability for analysis of data sets of any depth and coverage (Fig. 4, Table 1).

**Figure 4.**
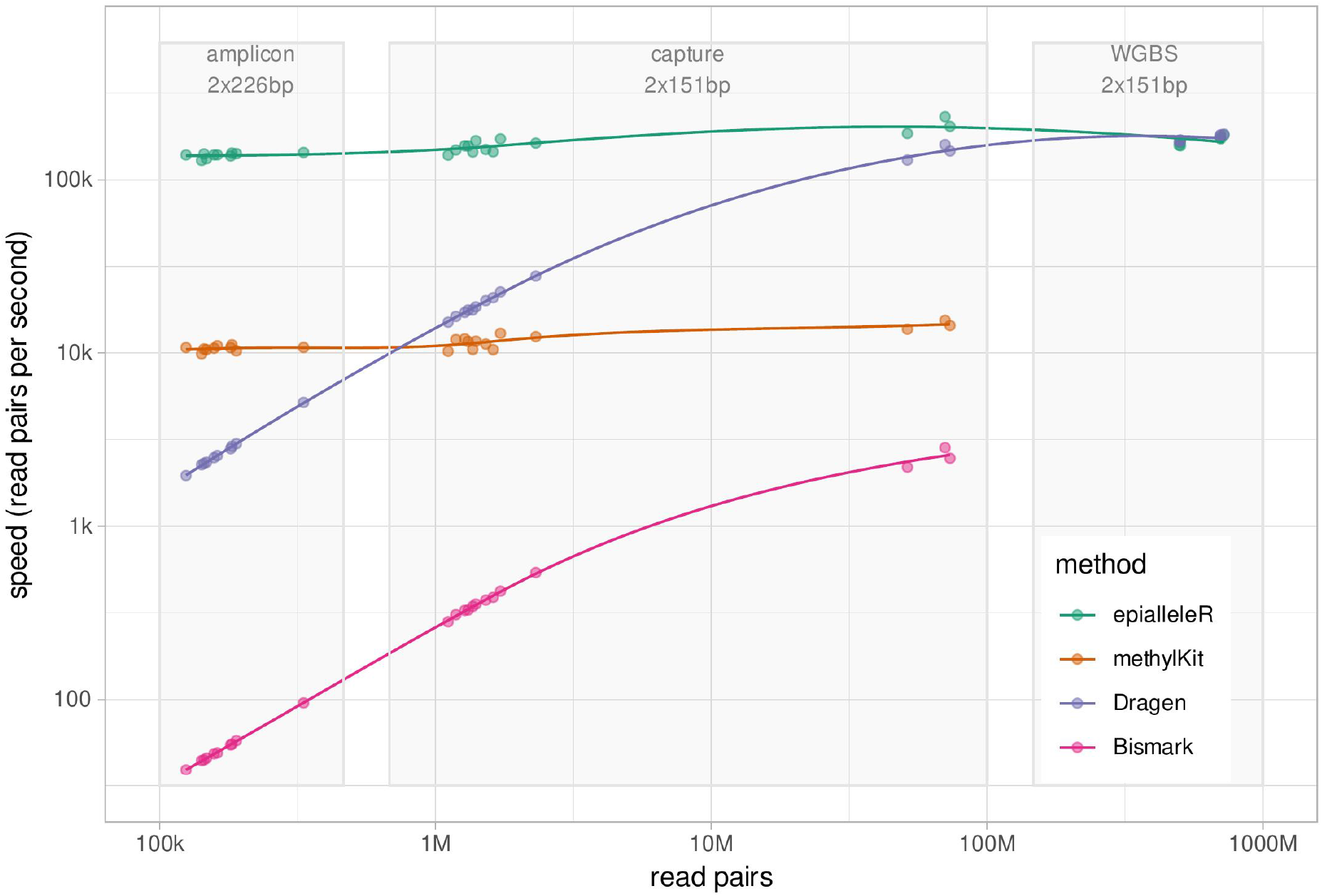
Data processing speed (in read pairs per second) of epialleleR as compared to three other methods for methylation reporting (methylKit, Bismark, and DRAGEN Bio-IT Platform). Read count (in number of pairs) is given at x-axis; light grey boxes outline data obtained by targeted amplicon-based, genome-wide capture-based, or whole-genome bisulfite sequencing.

## DISCUSSION

While conflicting data has linked low-level mosaic primary constitutional epimutations to cancer risk for more than a decade (Lønning et al. 2019), we have recently obtained firm evidence implicating primary epimutations within the *BRCA1* gene in an elevated risk of incident breast and ovarian cancer (Lønning et al. 2022). The assumption that such epimutations may affect other tumour suppressor genes and, therefore, lead to other cancer forms (Lønning et al. 2019), institutes a new research area with respect to cancer risk. Further, the findings of such epimutations in umbilical cord blood (Lønning et al. 2018) indicate prenatal events of a yet unknown genesis. This creates the need for multidisciplinary studies on the mechanisms of these events and on their effects in respect to cancer risk, as well as the need for ultrasensitive methods allowing sample assessment at a high scale.

Here, we present the details on a fast, accurate, and sensitive method to detect, quantify and visualise epialleles in NGS data. The method shows its superiority versus conventional methods of methylation reporting, especially when applied for detection of low-frequency methylation events, as it is by design less susceptible to variations in conversion efficiency or sequencing quality. Although epialleleR is not a differential methylation analysis tool, its output can be directly used to group samples based on their methylation profiles (by applying a simple threshold or using unsupervised clustering), as well as an input for other differential/aberrant methylation analysis software (the latter is not possible for WSH analysis tools). We thoroughly tested epialleleR using bisulfite sequencing data; the method, however, can also be applied to analyse and compare data obtained using any methylation sequencing technique (reduced representation bisulfite sequencing, RRBS; oxidative bisulfite sequencing, oxBS-Seq; Tet-assistant bisulfite sequencing, TAB-Seq), as long as methylation in these data can be called at individual cytosine residues instead of being analysed by comparing relative abundance of the fragments (such as for methylation sensitive restriction enzyme sequencing, MRE-Seq, or methylated DNA immunoprecipitation sequencing, MeDIP-Seq). Subtle though noticeable changes in cytosine reporting accuracy together with immense speed gain make epialleleR a method of choice not only for discovery of infrequent hypermethylated epialleles (as in (Lønning et al. 2022)), but also as a tool to produce conventional (no read thresholding) cytosine reports from any methylation sequencing alignment files. The implemented method is fully documented and can be easily used from within the R environment for statistical computing. With the epialleleR already revealing its suitability for detection of low-level mosaic methylation events in a large data set (Lønning et al. 2022), we believe it constitutes an optimal tool for assessment of low-level mosaic methylation with respect to risk of cancer as well as other diseases of relevance.

## CONCLUSIONS

Here, we present epialleleR, very fast, accurate, and sensitive method to detect, quantify and visualise epialleles in NGS data. Efficient implementation and improvements in cytosine reporting accuracy allow us to recommend epialleleR not only for analysis of methylation patterns and to enhance low-level differentially methylated region discovery, but also as a conventional cytosine reporting tool for various kinds of methylation sequencing data. The epialleleR R/Bioconductor package is freely available at https://bioconductor.org/packages/epialleleR/ and https://github.com/BBCG/epialleleR.

## MATERIALS AND METHODS

### Next-generation sequencing

White blood cells (WBC) DNA samples from anonymized males (n=88) (Knappskog et al. 2011, 2012) and human HCT116 DKO methylated DNA control sample (Zymo Research, cat.no. D5014-2) were bisulfite converted, and five DNA fragments, representing promoter regions of five established tumour suppressor genes, were amplified using custom set of primers (GRCh38 assembly coordinates of assayed regions: *MLH1*, chr3:36993123–36993500; *CDKN2A*, chr9:21974554–21974921; *MGMT*, chr10:129467118–129467477; *CDH1*, chr16:68737102–68737469; *BRCA1*, chr17:43125171– 43125550), indexed, and sequenced similarly to as previously described (Lønning et al. 2022) (GSE201688). The resulting average coverage was 5000x–50000x per amplicon.

### Bioinformatic and statistical analyses

Massive parallel sequencing (NGS) reads were mapped/aligned to the GRCh38 human reference genome and the methylation was called using Illumina DRAGEN Bio-IT Platform (v3.9.5) with the following parameters: --methylation-mapping-implementation single-pass, --enable-methylation-calling true, --methylation-generate-cytosine-report false, --methylation-protocol non-directional, -- enable-sort false, unless stated otherwise. R software environment for statistical computing (v4.1.2) was used for all downstream statistical analyses.

The frequency of hypermethylated alleles across assayed regions in n=88 male WBC DNA samples were estimated using epialleleR::generateAmpliconReport with the following parameters: min.mapq=30, min.baseq=20, nthreads=4, threshold.reads=TRUE, report.context=“CG”, and bed.file pointing to a location of BED (browser extensible data) file with genomic regions amplified (see amplicon coordinates above). Two sample subgroups (n=8 and n=10) were selected for sensitivity analyses based on the frequencies of hypermethylated alleles as explained below.

### Cytosine reporting accuracy comparison

Four sets of paired-end sequencing reads (151bp, 50 million read pairs each set) were simulated using Sherman Bisulfite FastQ Read Simulator (Krueger 2011) with the following options: --length 151, -- number_of_seqs 50000000, --paired_end, --minfrag 70, --maxfrag 400, --CG_conversion 0, -- CH_conversion 99.5 and varying sequencing error rate (−-error_rate parameter) of 0%, 0.1%, 0.3% or 0.6%. Human chromosome 19 sequence (GRCh38.p13 NC_000019.10, 58617616 bp, 1105620 forward strand CpGs) was used as a reference genome for read simulation and mapping/alignment due to its highest CpG content across all human chromosomes (Harris et al. 2020) and in order to maintain optimal balance of analysis speed and base coverage. Each set of reads was then duplicated, all read1 cytosines (C) and read2 guanines (G) in any context in the duplicate sets were replaced with thymines (T) and adenines (A), respectively. Then, duplicate sets (i.e., unmethylated reads) were merged with original sets (i.e., methylated reads) resulting in four sets of reads 100 million pairs each, with the cytosine conversion rate of exactly 50% and about 99.75% in CpG and non-CpG contexts, respectively.

The mapping and alignment of simulated reads was performed using Illumina DRAGEN Bio-IT Platform v3.9.5 with the following modification in parameters: --methylation-protocol directional. Methylation reporting was done as described below (Speed comparison section).

### Sensitivity comparison on admixed samples

In order to simulate variable methylation levels while maintaining biological heterogeneity of the samples, we selected ten male DNA NGS samples with the lowest frequency of hypermethylated alleles across all assayed regions, then admixed varying fractions of reads from two random samples and additionally “spiked” certain number of fully methylated reads from methylated DNA control sample. This resulted in 150 samples containing 0%, 0.01%, 0.03%, 0.1%, 0.3% or 1% of methylated reads per sample (25 samples per every category).

Read mapping, alignment, methylation calling, and generation of genome-wide cytosine reports was performed using Illumina DRAGEN Bio-IT Platform as described above. VEF calling was performed using epialleleR::generateCytosineReport with the following parameters: min.mapq=0, min.baseq=0, nthreads=4, threshold.reads=TRUE, report.context=“CG”.

Methylation patterns and per-read beta values for all samples were extracted using epialleleR::extractPatterns with the following parameters: min.mapq=30, min.baseq=20, nthreads=4, clip.patterns=FALSE, and bed.file pointing to a location of BED file with genomic regions amplified (see amplicon coordinates above).

Barnes-Hut t-Distributed Stochastic Neighbor Embedding (t-SNE) analysis was performed using R package Rtsne v0.15 (Krijthe 2015) and matrices of beta or VEF values for all genomic positions of CpGs with the coverage of at least 1000x and available values for all analysed samples (total number of CpGs, n=138; *MLH1*, n=20; *CDKN2A*, n=35; *MGMT*, n=33; *CDH1*, n=32; *BRCA1*, n=18).

### Sensitivity comparison to methylation array data

Eight additional WBC DNA NGS samples from anonymized males carrying hypermethylated alleles in at least one of the assayed regions were selected, and VEF calling was performed using epialleleR::generateCytosineReport with the following parameters: min.mapq=30, min.baseq=20, nthreads=4, threshold.reads=TRUE, report.context=“CG”. The same DNA samples were also bisulfite converted using the Zymo EZ DNA Methylation Kit (Zymo Research, cat.no. D5001), and genome-wide methylation levels were assessed using Illumina HumanMethylationEPIC BeadChip arrays according to manufacturer’s instructions. Resulting IDAT files were processed (normalized and annotated) with the minfi Bioconductor package (Aryee et al. 2014) using the preprocessQuantile method with outlier thresholding enabled (GSE201689). For direct comparison, only the CpGs that are covered in all samples by both BeadChip arrays (p-value of 0) and targeted sequencing (minimum sequencing coverage of 5000x) were retained (*MLH1*, n=10; *CDKN2A*, n=2; *MGMT*, n=4; *CDH1*, n=7; *BRCA1*, n=14). Pairwise sample comparison was performed using t-test with Holm adjustment for multiple comparisons.

The sets of CpGs that are differentially methylated between cell blood types were reported previously: DNA methylation profiles for six blood cell types from six males (Reinius et al. 2012; Jaffe 2022), and DNA methylation profiles for seven blood cell types from cord blood of 104 newborns (Bakulski et al. 2016; Andrews and Bakulski 2022). CpG-level differential methylation analysis p values were Holm-adjusted and the ones remained significant (adjusted p≤0.05; n=73629 of total 456655 for male blood data set; n=221246 of total 429794 for newborn cord blood data set) were checked for overlap with the set of CpGs analysed in this study (n=35 CpGs of total n=37 were present in each of male/newborn data sets).

### Differential methylation analysis

Differentially methylated regions (DMRs) were called using R package DMRcate (v2.12.0) with the following parameters: lambda=1000, min.cpgs=2, pcutoff=“fdr” (Peters et al. 2021). Aberrantly methylated regions were called using R package ramr (v1.6.0) with the following parameters: ramr.method=“beta”, min.cpgs=2, merge.window=500 (Nikolaienko et al. 2022). To enable maximum likelihood estimation of beta distribution parameters, all zeros were replaced with minimum double values (2.26e-308).

For intergroup DMR discovery in admixed samples, pairwise comparison of sample groups defined by the amount of admixed reads was performed (n=25 samples in each group) using the default level of false discovery rate (FDR) cutoff (equals 0.05). For DMR discovery in real samples, as DMRcate methods require two classes/categories for comparison, every real sample from the test dataset was tested against all the other samples using the default FDR cutoff value.

To assess DMR (or AMR) recall metrics in admixed samples, every sample with admixed reads was compared using DMRcate (or ramr) to the group of 25 samples without admixed reads at a varying level of FDR (or p value) cutoff of 0.05, 0.01, or 0.001. As the admixed reads covered all five assayed regions, only the total number of real positive regions (P, equals 5 for each comparison), the number of true positive regions (TP), and the number of false negative regions (FN=P–TP) were known, while the numbers of true negative (TN) or false positive (FP) regions were undefined. Therefore, recall, or true positive rate (TPR=TP/P) was chosen as a sensitivity metric.

### Within-sample heterogeneity

Estimation of within-sample heterogeneity (WSH) was performed on eight samples used in the sensitivity comparison between array- and NGS-based methylation profiling. Difference in entropy was evaluated using methclone (v1) (Li et al. 2014) with a distance cutoff of 500 and minimum read coverage of 1000 for every pair of samples. As methclone outputs values for multiple genomic regions, the minimum value (representing absolute largest difference) was selected and used further. Entropy, epipolymorphism, fraction of discordant read pairs (FDRP) and proportion of discordant reads (PDR) were evaluated using R package WSH (v0.1.6) (Scherer et al. 2020) with the following options: mapq.filter=30, window.size=500, and bam.file pointing to a location of BAM file. Due to exponential complexity of FDRP calculation, option max.reads was set to 100 for FDRP calculation and to 1e+06 otherwise. Pairwise sample score comparison was performed using t-test with Holm adjustment for multiple comparisons.

### Speed comparison

Comparison of processing speed was performed on 29 BAM files containing paired-end alignments and methylation calls derived from bisulfite sequencing of human WBC DNA samples prepared using the following assays: A) amplicon-based sequencing of promoter regions of *BRCA1* gene (n=10 files, 0.12–0.33 million read pairs per file, average coverage of ∼20,000x) (Lønning et al. 2022); B) genome-wide capture-based bisulfite sequencing of promoter regions of 283 tumour suppressor genes (n=10 files, 1.11–2.31 million read pairs per file, average coverage of ∼60x; and n=3 files, 51.4–73.4 million read pairs per file, average coverage of ∼1000x) (Poduval et al. 2020; Eikesdal et al. 2021); C) whole-genome bisulfite sequencing (n=6 files, 497–723 million read pairs per file, average coverage of ∼60x; epialleleR and Illumina DRAGEN Bio-IT Platform only). The two former data sets (A and B) were generated in-house and described previously, while the latter data (C) were obtained from NCBI Sequence Read Archive (GEO/SRA samples GSM3683953/SRX6640720, GSM3683958/SRX6640725, GSM3683965/SRX6640732, GSM3683951/SRX6640718, GSM3683955/SRX6640722, and GSM3683962/SRX6640729) and reported elsewhere (Zhou et al. 2019).

Processing times to produce conventional cytosine reports were recorded as following: Bismark CX methylation reports were created using Bismark v0.22.3 (Krueger and Andrews 2011) with the following parameters: command bismark_methylation_extractor, --paired-end, --no_overlap, --comprehensive, --gzip, --mbias_off --parallel 8, --cytosine_report, --CX, --buffer_size 64G. As parallel processing was requested, Bismark used up to 24 cores for some of its subtasks.

methylKit CX methylation reports were created using R/Bioconductor package methylKit v1.20.0 (Akalin et al. 2012) with the following parameters: function methylKit::processBismarkAln, minqual=0, mincov=0,save.context=c(“CpG”,”CHG”,”CHH”), nolap=TRUE and location pointing to the location of a BAM file. Parallel processing is currently not available for methylKit::processBismarkAln.

epialleleR CX methylation reports were created using R/Bioconductor package epialleleR v1.3.5 with the following parameters: function epialleleR::generateCytosineReport, min.mapq=0, min.baseq=0, nthreads=4 (number of HTSlib decompression threads), threshold.reads=FALSE, report.context=“CX” and bam pointing to the location of a BAM file. epialleleR methods currently run in a single-threaded mode only but can benefit from additional BAM decompression threads provided by HTSlib.

Bismark, methylKit and epialleleR were tested on the workstation equipped with AMD EPYC 7742 64-core processor, 512GB of memory and the Red Hat Enterprise Linux Server release 7.9 (Developer Toolset 6, GCC v6.3.1), with BAM files retrieved from high-speed (10Gbps) network accessible storage.

DRAGEN CX methylation reports were created using Illumina DRAGEN Bio-IT Platform v3.9.5 (Intel Xeon Gold 6126 48-core processor, 256GB of memory and CentOS Linux release 7.5.1804) with the following parameters: --methylation-generate-cytosine-reports=true, --enable-sort=false, --enable-duplicate-marking=false, --methylation-report-only=true and --bam-input pointing to the location of a BAM file. Default number of threads (up to 24) were used for data processing using DRAGEN; BAM files were accessed from local, high-speed NVMe solid state disk.

For Bismark and DRAGEN, elapsed time measurements were stably reproducible, thus processing time was recorded only once for each file. For methylKit and epialleleR, the tests were run five times in sequential random order by means of R package microbenchmark v1.4.9, and the average time was used in comparison to mitigate variability in processing time measurements.

## DECLARATIONS

### Ethics approval and consent to participate

Ethics approvals and other relevant information for patient-generated data used in speed assessment were included and described in previous studies (Poduval et al. 2020; Eikesdal et al. 2021; Lønning et al. 2022; Knappskog et al. 2011). All analyses of biomaterial were approved by Regional Ethics Committees for medical research and all samples were collected after written informed consent from the sample donors (REK-vest Norway reference numbers: 3.2008.1932, 2015/1493 and 2018/1566.

### Availability of data and materials

The epialleleR R/Bioconductor package is freely available at https://bioconductor.org/packages/epialleleR/ and https://github.com/BBCG/epialleleR. The R scripts used in this manuscript and the data underlying accuracy and sensitivity analyses are freely available at DataverseNO (https://doi.org/10.18710/2BQTJP). Sensitive data used for the processing speed assessment are available from the authors in accordance with study protocols.

Public data for sensitivity analysis have been deposited at NCBI Gene Expression Omnibus under accession number GSE201690. Public whole-genome bisulfite sequencing data used for the processing speed assessment are available at NCBI Sequencing Read Archive under accession number SRP217135.

Supplementary Data are available online.

### Authors contribution

Conceived the project: O.N., P.E.L., S.K. Supervised the project: P.E.L., S.K. Conceived, designed, and implemented the software and the analysis pipeline: O.N. Wrote the paper: O.N., P.E.L., S.K. All authors read and approved the final manuscript.

