## Supplementary Material for "epialleleR: an R/Bioconductor package for sensitive allele-specific methylation analysis in NGS data"

Supplementary Table 1. Complete metrics of cytosine reporting across all three possible cytosine genomic contexts (CHH, CHG and CpG), obtained using simulated chr19 reads with varying sequencing error rate by selected tools. “reported”, number of cytosines present in the cytosine reports; “valid context”, number of cytosines for which genomic context was correctly identified; “invalid context”, number of cytosines for which genomic context was incorrectly identified; “not covered”, number of cytosines not present in a cytosine report; “mean coverage”, average coverage of all cytosines in this context; “mean” and “variance”, average value and variance for beta values of all cytosines in this context; “is 0.5”, number of cytosines with beta value of exactly 0.5 (ground truth for this dataset); “is not 0.5”, number cytosines with beta value not equal to 0.5.

Supplementary Fig. 1. Scaled density of per-read beta values from all admixed samples combined, split by the level of admixed reads and genomic region of interest. The y axis is scaled to 1 and limited to 0.015. The hypermethylated ($\beta \geq 0.5$) reads are increasingly apparent on the right sides of plots in accordance with an increase in admixed methylated reads.

Supplementary Fig. 2. Methylation patterns from all admixed samples combined, split by the level of admixed reads and genomic region of interest. Lines depict patterns, open and closed circles depict unmethylated and methylated cytosines, respectively. Numbers on the right of every pattern indicate how many times each pattern occurs for every given gene/sample set. Due to very high number of methylation pattern types, only the most abundant pattern (if any) is shown for each range of average beta value: [0,0.2), [0.2,0.4), [0.4,0.6), [0.6,0.8), [0.8,1]. The hypermethylated ($\beta \geq 0.5$) patterns are increasingly apparent at the top of plots in accordance with an increase in admixed methylated reads.

Supplementary Fig. 3. Scaled density of per-read beta values from n=8 real samples used to compare sensitivity of methylation profiling by NGS and array, split by sample and genomic region of interest. The y axis is scaled to 1 and limited to 0.015. The population of hypermethylated ($\beta \geq 0.5$) reads that are apparent on the Fig. 3A in the main text are pointed to by black arrows.

Supplementary Fig. 4. Methylation patterns from n=8 real samples used to compare sensitivity of methylation profiling by NGS and array, split by sample and genomic region of interest. Lines depict patterns, open and closed circles depict unmethylated and methylated cytosines, respectively. Numbers on the right of every pattern indicate how many times each pattern occurs for every given gene/sample set. Due to very high number of methylation pattern types, only the most abundant pattern (if any) is shown for each range of average beta value: [0,0.2), [0.2,0.4), [0.4,0.6), [0.6,0.8), [0.8,1]. The hypermethylated ($\beta \geq 0.5$) patterns represent hypermethylated epialleles that are present in certain samples/regions, as shown by the Fig. 3A in the main text and Supplementary Fig. 3.

Supplementary Fig. 5. (A) Distribution of per-read beta values for NGS reads covering CpGs that are common for NGS and BeadChip array. For clarity, only the reads with average beta of at least 0.5 (i.e., representing hypermethylated epialleles) are included. Single observations are shown as dots, number of observations is given above. (B) Lower panel: box plots of NGS-derived VEF values for individual CpGs; upper panel: significant aberrantly or differentially methylated regions identified by ramr or DMRcate, respectively, based on VEF values. (C) Lower panel: box plots of NGS-derived beta values for individual CpGs; upper panel: significant aberrantly or differentially methylated regions identified by ramr or DMRcate, respectively, based on NGS-derived beta values. (B–C) The lower and upper hinges of boxes correspond to the first (Q1) and third (Q3) quartiles; the bar in the middle correspond to the median value; the upper and lower whisker extend to Q3+1.5*IQR and Q1-1.5*IQR, respectively, while the values outside this range (outliers) are plotted as dots. Zero values are not plotted. The colouring is preserved for n=8 samples used in Fig. 3 of the main text. The n=10 samples used to create admixed samples and not included in Fig. 3 are plotted in light grey.

Supplementary Fig. 6. (A) Heatmap of minimum (equals absolute largest) difference in combinatorial entropy for all pairs of samples, split by genomic region of interest. (B) Heatmap of combinatorial entropy, epipolymorphism, fraction of discordant read pairs (FDRP) and proportion of discordant reads (PDR), split by genomic region of interest. (C) Heatmap of p values for pairwise comparison of samples using WSH scores, split by score and genomic region of interest. *** $p<0.001$, ** $p<0.01$, * $p<0.05$, blank $p \geq 0.05$

Supplementary Table 1

| sequencing error rate | method | CHH | | | | | | CHG | | | | | | CpG | | | | | | | |
|---|---|---|---|---|---|---|---|---|---|---|---|---|---|---|---|---|---|---|---|---|---|
| | | reported | | not covered | mean coverage | beta values | | reported | | not covered | mean coverage | beta values | | reported | | not covered | mean coverage | beta values | | | |
| | | valid context | invalid context | | | mean | variance | valid context | invalid context | | | mean | variance | valid context | invalid context | | | is 0.5 | is not 0.5 | mean | variance |
| 0.00% | dragen | 19154385 | 0 | 56187 | 203.68 | **0.002501** | 1.28E-05 | 6573937 | 0 | 19963 | 203.80 | **0.002500** | 1.27E-05 | 2200279 | 0 | 10961 | 203.25 | 2189866 | 10413 | 0.499997 | 5.96E-07 |
| | bismark | 19154371 | 0 | 56201 | 194.96 | 0.002501 | 1.34E-05 | 6573935 | 0 | 19965 | 195.07 | 0.002500 | 1.33E-05 | 2200275 | 0 | 10965 | 194.56 | **2191308** | **8967** | **0.499997** | **5.92E-07** |
| | methylkit | 19154385 | 0 | 56187 | 204.07 | 0.002503 | **1.28E-05** | 6573937 | 0 | 19963 | 204.19 | 0.002501 | **1.27E-05** | 2200279 | 0 | 10961 | 203.64 | 2189867 | 10412 | 0.499997 | 5.97E-07 |
| | epialleleR | 19154385 | 0 | 56187 | 203.68 | **0.002501** | 1.28E-05 | 6573937 | 0 | 19963 | 203.80 | **0.002500** | 1.27E-05 | 2200279 | 0 | 10961 | 203.25 | 2189866 | 10413 | 0.499997 | 5.96E-07 |
| 0.10% | dragen | 19154394 | 0 | 56178 | 203.54 | 0.002661 | 1.37E-05 | 6573909 | 0 | 19991 | 203.66 | 0.002662 | 1.36E-05 | 2200248 | 0 | 10992 | 203.17 | 2116830 | 83418 | 0.499835 | 1.73E-06 |
| | bismark | 19154378 | 0 | 56194 | 194.83 | 0.002646 | 1.42E-05 | 6573904 | 0 | 19996 | 194.95 | 0.002648 | 1.41E-05 | 2200248 | 0 | 10992 | 194.49 | 2127434 | 72814 | 0.499850 | 1.69E-06 |
| | methylkit | 19154394 | 156 | 56178 | 203.92 | 0.002662 | 1.37E-05 | 6573909 | 116 | 19991 | 204.05 | 0.002664 | 1.36E-05 | 2200248 | 107 | 10992 | 203.55 | 2116536 | 83712 | 0.499835 | 1.75E-06 |
| | epialleleR | 19154393 | 0 | 56179 | 203.57 | **0.002624** | **1.35E-05** | 6573909 | 0 | 19991 | 203.69 | **0.002626** | **1.34E-05** | 2200248 | 0 | 10992 | 203.20 | **2132685** | **67563** | **0.499872** | **1.54E-06** |
| 0.30% | dragen | 19154462 | 0 | 56110 | 203.24 | 0.002977 | 1.53E-05 | 6573961 | 0 | 19939 | 203.34 | 0.002980 | 1.53E-05 | 2200275 | 0 | 10965 | 202.80 | 1978460 | 221815 | 0.499504 | 3.22E-06 |
| | bismark | 19154457 | 0 | 56115 | 194.58 | 0.002928 | 1.57E-05 | 6573957 | 0 | 19943 | 194.67 | 0.002931 | 1.57E-05 | 2200273 | 0 | 10967 | 194.16 | 2007647 | 192626 | 0.499554 | 3.03E-06 |
| | methylkit | 19154462 | 1849 | 56110 | 203.63 | 0.002978 | 1.52E-05 | 6573961 | 1639 | 19939 | 203.72 | 0.002982 | 1.52E-05 | 2200275 | 1547 | 10965 | 203.18 | 1978911 | 221364 | 0.499506 | 3.21E-06 |
| | epialleleR | 19154459 | 0 | 56113 | 203.34 | **0.002857** | **1.47E-05** | 6573960 | 0 | 19940 | 203.43 | **0.002860** | **1.47E-05** | 2200275 | 0 | 10965 | 202.90 | **2028765** | **171510** | **0.499628** | **2.59E-06** |
| 0.60% | dragen | 19154927 | 0 | 55645 | 202.79 | 0.003498 | 1.79E-05 | 6573979 | 0 | 19921 | 202.90 | 0.003506 | 1.78E-05 | 2200281 | 0 | 10959 | 202.36 | 1763714 | 436567 | 0.498942 | 1.29E-05 |
| | bismark | 19154901 | 0 | 55671 | 194.19 | 0.003393 | 1.81E-05 | 6573971 | 0 | 19929 | 194.30 | 0.003402 | 1.81E-05 | 2200279 | 0 | 10961 | 193.79 | 1819202 | 381077 | 0.499051 | 1.25E-05 |
| | methylkit | 19154927 | 9913 | 55645 | 203.17 | 0.003497 | 1.78E-05 | 6573979 | 8730 | 19921 | 203.28 | 0.003501 | 1.78E-05 | 2200281 | 8115 | 10959 | 202.74 | 1767002 | 433279 | 0.498952 | 1.28E-05 |
| | epialleleR | 19154923 | 0 | 55649 | 202.99 | **0.003237** | **1.66E-05** | 6573976 | 0 | 19924 | 203.11 | **0.003241** | **1.65E-05** | 2200279 | 0 | 10961 | 202.56 | **1866484** | **333795** | **0.499222** | **1.14E-05** |

Supplementary Figure 1. Scaled density of per-read beta values in admixed samples

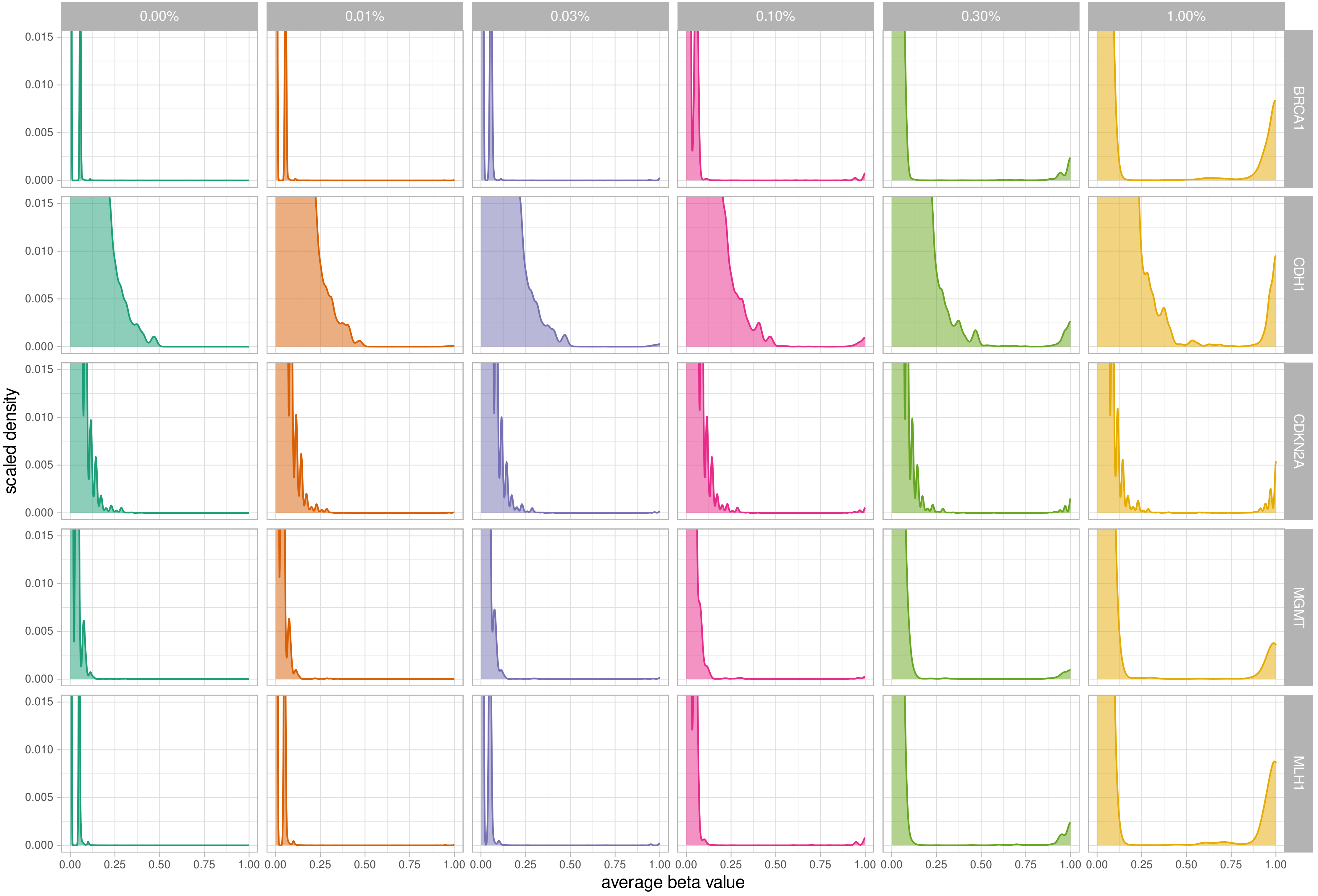

Supplementary Figure 2. Epialleles in admixed samples

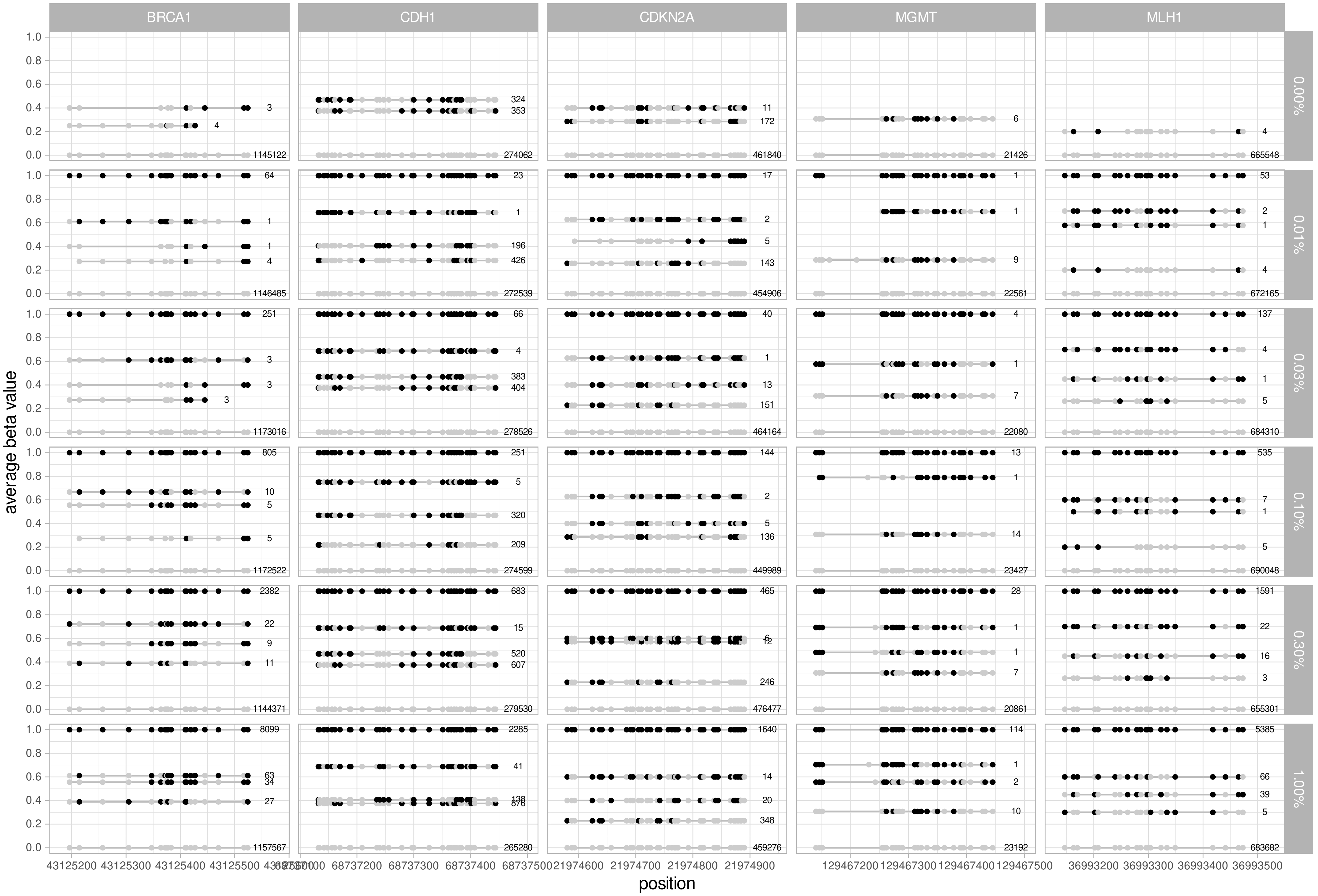

Supplementary Figure 3. Scaled density of per-read beta values in real samples

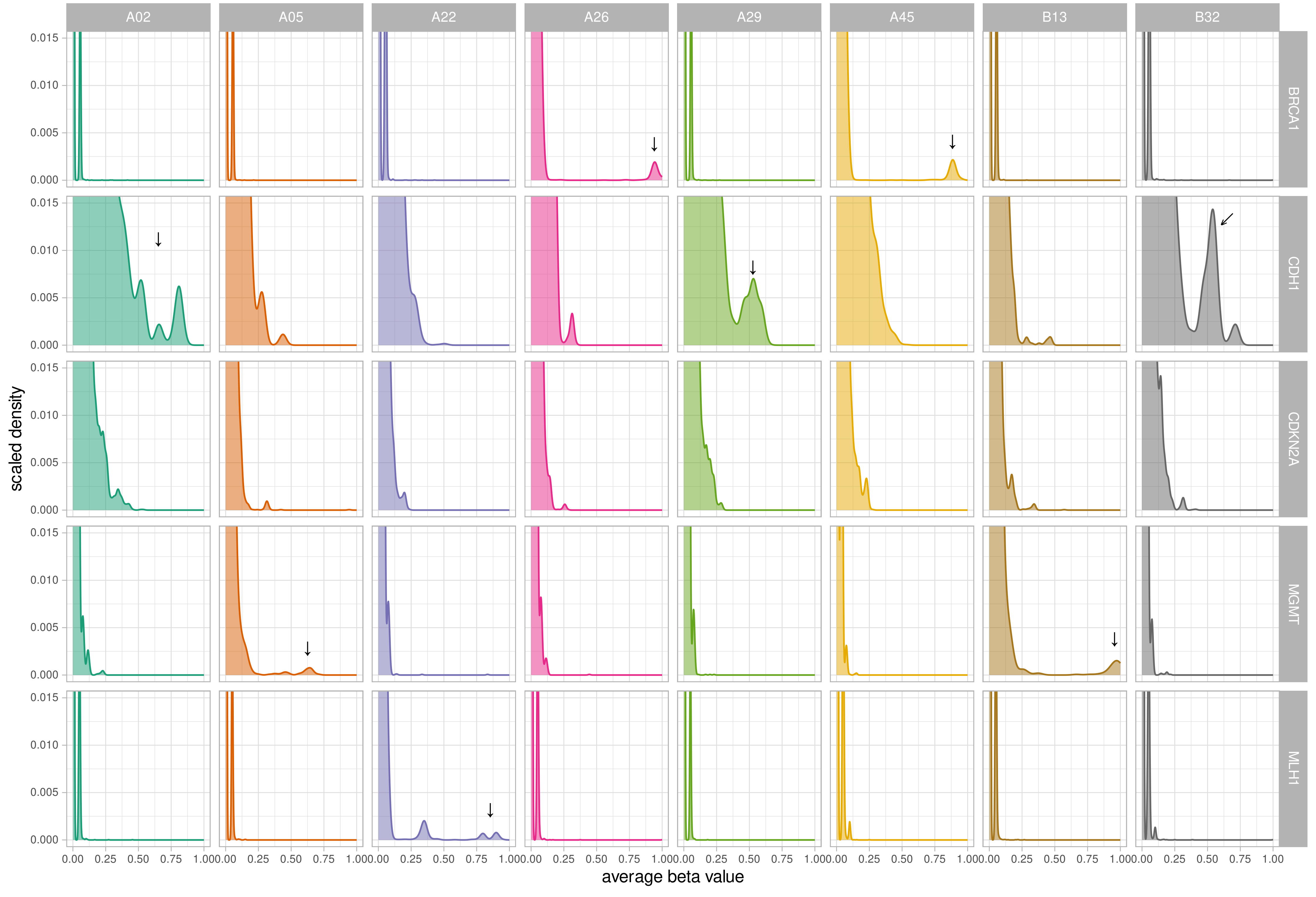

### Supplementary Figure 4. Epialleles in real samples

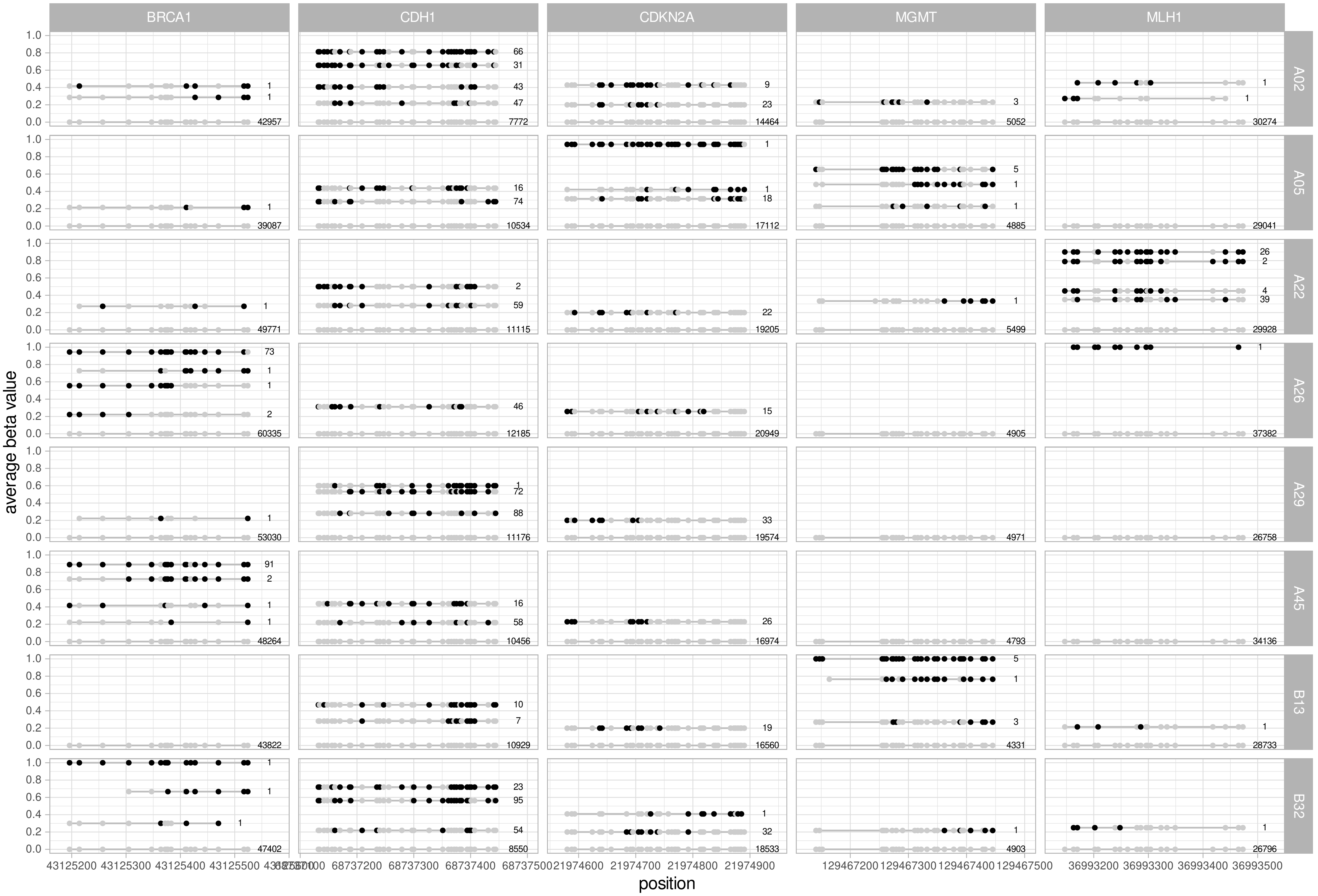

Supplementary Figure 5

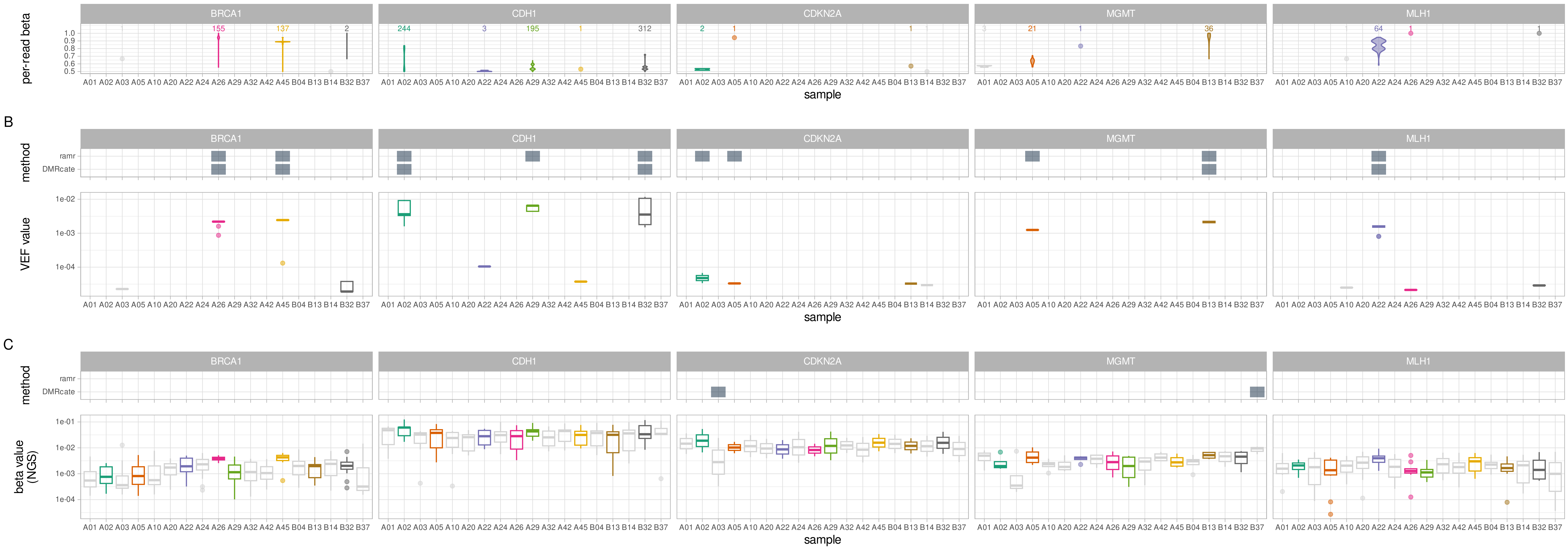

Supplementary Figure 6

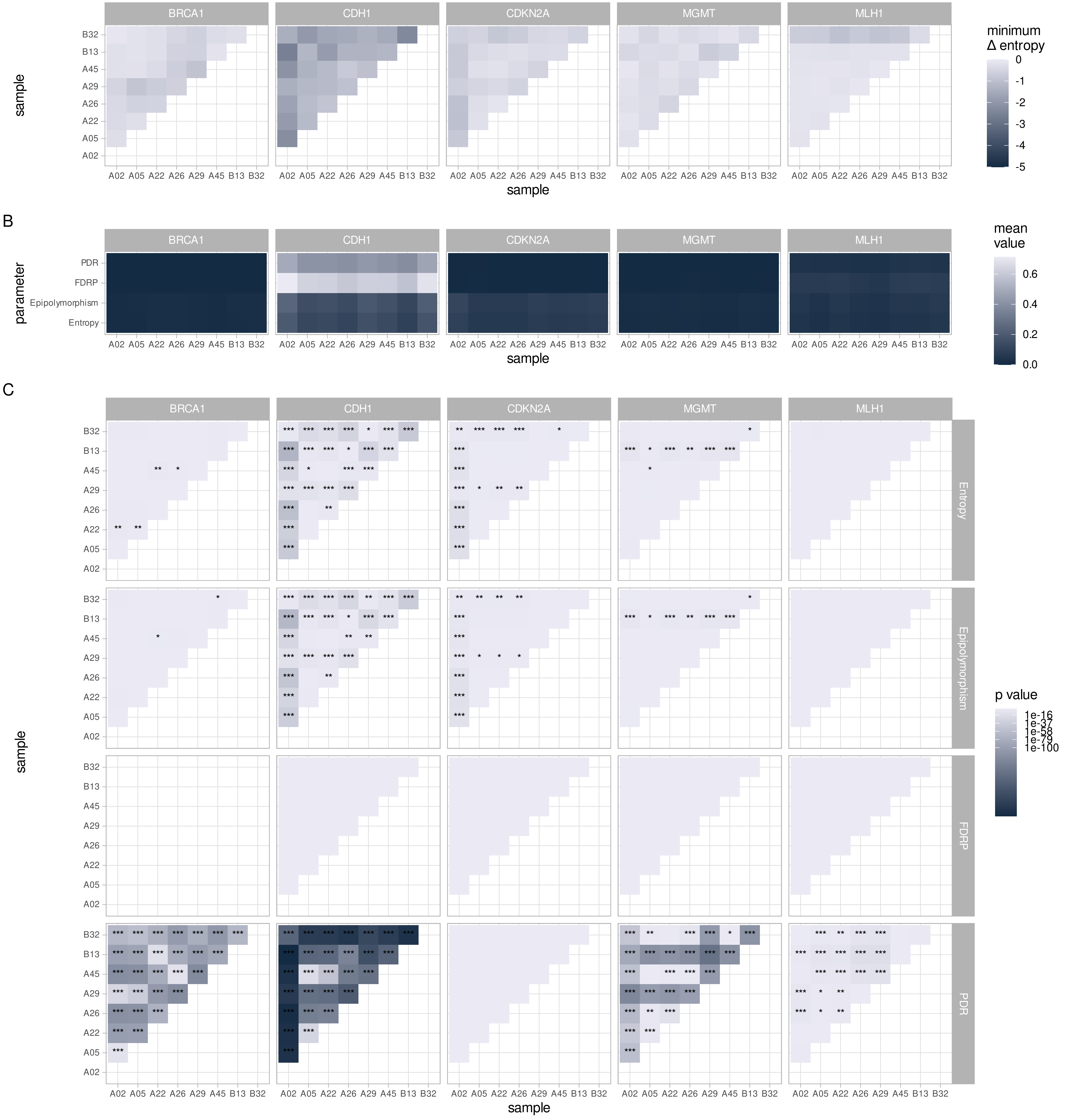
